# Soundscapes predict species occurrence in tropical forests

**DOI:** 10.1101/2020.09.24.311381

**Authors:** Sarab S. Sethi, Robert M. Ewers, Nick S. Jones, Jani Sleutel, Adi Shabrani, Nursyamin Zulkifli, Lorenzo Picinali

## Abstract

1. Accurate occurrence data is necessary for the conservation of keystone or endangered species, but acquiring it is usually slow, laborious, and costly. Automated acoustic monitoring offers a scalable alternative to manual surveys, but identifying species vocalisations requires large manually annotated training datasets, and is not always possible (e.g., for silent species). A new, intermediate approach is needed that rapidly predicts species occurrence without requiring extensive labelled data.
2. We investigated whether local soundscapes could be used to infer the presence of 32 avifaunal and seven herpetofaunal species across a tropical forest degradation gradient in Sabah, Malaysia. We developed a machine-learning based approach to characterise species indicative soundscapes, training our models on a coarsely labelled manual point-count dataset.
3. Soundscapes successfully predicted the occurrence of 34 out of the 39 species across the two taxonomic groups, with area under the curve (AUC) metrics of up to 0.87 (Bold-striped Tit-babbler *Macronus bornensis)*. The highest accuracies were achieved for common species with strong temporal occurrence patterns.
4. Soundscapes were a better predictor of species occurrence than above-ground biomass – a metric often used to quantify habitat quality across forest degradation gradients.
5. *Synthesis and applications*: Our results demonstrate that soundscapes can be used to efficiently predict the occurrence of a wide variety of species. This provides a new direction for audio data to deliver large-scale, accurate assessments of habitat suitability using cheap and easily obtained field datasets.

## Introduction

Ecosystems are being subjected to increasing external pressures from land-use change and global warming (Lambin and Meyfroidt, 2011; Walther et al., 2002). These pressures have resulted in global biodiversity declines, as the natural habitats required to support many species shrink and disappear (Newbold et al., 2015). Efforts to slow this decline often aim to protect areas of high conservation value that may support populations of endangered or keystone species (Mills et al., 1993). This leads to the key question; how can we identify such locations rapidly, accurately, and on a large scale?

An established solution is to carry out manual surveys of the region of interest (Brown et al., 2013). Common approaches include actively searching for species of interest, deploying traps to capture them, or looking for features that may indicate their presence (e.g., nests). However, manual surveys are expensive, labour intensive, and do not scale well temporally or spatially (Gijzen, 2013). In contrast, automated acoustic monitoring has shown promise as a route to gaining scalable insight into ecological systems (Gibb et al., 2019). Audio data can be recorded and analysed inexpensively, in real-time, and over extended durations, making it an increasingly powerful tool for ecologists and conservationists (Hill et al., 2018; Sethi et al., 2018; Sethi et al., 2020a).

Species occurrence data can be extracted from audio recordings automatically by detecting vocalisations. Using a large training dataset of annotated examples, a machine learning model can learn to identify calls made by a target species (Clink et al., 2019; Stowell et al., 2016; Wrege et al., 2017). This approach, however, relies upon three key assumptions; (i) the species has a unique vocalisation, (ii) the species is active and audible during the recording, and (iii) there exists a large labelled dataset of the species’ vocalisations (or the resources to collate such training data from scratch). These barriers are particularly difficult to overcome when searching for rare or endangered species in highly biodiverse and noisy environments such as tropical forests (Gibb et al., 2019; Stowell et al., 2018), or for species that are largely silent.

Analysing soundscapes in their entirety provides an alternate route to the automated analysis of eco-acoustic data (Pijanowski et al., 2011). In this approach, features of the audio signal are used to directly infer habitat quality, without the need for species specific training data (Pieretti et al., 2011; Sethi et al., 2020b; Sueur et al., 2008). Whilst soundscape features have been shown to correlate with high-level metrics of biodiversity, they are not normally used to provide direct evidence for how suitable a habitat is for a given species.

In this study we demonstrate that an environment’s soundscape can in fact be used as a powerful indicator of species occurrence. Rather than focussing on species-specific vocalisations, our model learned acoustic features which indicated species presence using only coarsely-labelled point count data from across a gradient of tropical forest degradation in Sabah, Malaysia. We were able to predict occurrence accurately for a number of avifaunal and herpetofaunal species without the need for large, precisely annotated training datasets. Additionally, we showed that soundscapes are a more accurate predictor of species occurrence than above-ground biomass, a metric often used to quantify habitat quality across forest degradation gradients (Pfeifer et al., 2015). Our findings indicate a promising new route for audio data to be used for the conservation of species on a large scale, and across a wide range of taxa, without many of the limitations of vocalisation detection-based approaches.

## Materials and methods

### Study location and estimates of habitat quality

This work was undertaken at the Stability of Altered Forest Ecosystems (SAFE) Project in Sabah, Malaysia (Ewers et al., 2011) between March 2018 and February 2020. We surveyed eleven sites across a land-use intensity gradient: two sites in oil palm plantations, two sites in salvage logged forest (last logged in the early 2010’s), five sites in selectively twice-logged forest (logged in the 1970’s and early 2000’s), and two sites in forest inside a protected area (where small amounts of illegal logging activity had occurred).

From 2012 to 2013, Pfeifer and colleagues (Pfeifer et al., 2015) conducted ground surveys of over 100 vegetation plots (25 x 25 m) across the SAFE project landscape to quantify above ground biomass (AGB). We averaged AGB from all surveyed plots within 1 km of each of our sampling sites (mean plots per site = 8.5, range = 2-16), for use as a quantitative measure of habitat quality.

### Avifaunal and herpetofaunal point counts

Across the 11 sampling sites, we carried out 790 avifaunal and 771 herpetofaunal point counts (of which 483 were undertaken simultaneously). Each point count lasted 20 minutes and surveys were distributed evenly throughout the 24 hours of the day, giving approximately three replicates per site per hour for both avifaunal and herpetofaunal point counts.

During point counts, we recorded all visual or aural encounters of avifaunal or herpetofaunal species within a 10 m radius of the sampling site. Species were cross-referenced with the Global Biodiversity Information Facility (GBIF) backbone taxonomy to validate taxonomic classifications (GBIF Secretariat, 2020).

Occurrence data (presence/absence) was thus acquired for 175 avifaunal and 53 herpetofaunal species. Species present in fewer than 50 point counts were removed from the dataset. For those species classified as vulnerable or critically endangered by the IUCN Red List (Baillie et al., 2004), a reduced threshold of 15 occurrences was used. In total this gave us a set of 32 avifaunal and seven herpetofaunal species (Supp. Table S1). Five of the 32 avifaunal species were listed as vulnerable or critically endangered, but none of the seven herpetofaunal species were.

### Audio data and acoustic feature extraction

During each point count a simultaneous 20-minute audio recording was made using a Tascam DR-05 recorder mounted at chest height. Raw audio data was recorded to a single channel at 44.1 kHz in the WAV format.

We calculated learned acoustic features of the audio using a pretrained convolutional neural network (CNN), “VGGish”, developed by Hershey et. al (Hershey et al., 2017). The CNN was trained to perform a general-purpose audio classification task using an extremely large annotated dataset (Gemmeke et al., 2017), resulting in a general 128-dimensional acoustic feature embedding. Prior work has shown that embedding eco-acoustic data using this approach allows multi-scale monitoring of ecosystems and efficient characterisation of soundscapes (Sethi et al., 2020b).

The VGGish CNN takes a 16 kHz log-scaled Mel-frequency spectrogram as an input (96 temporal frames, 64 frequency bands) providing one feature vector per 0.96 s of audio. Since our raw audio data was recorded at a higher sample rate, we pre-processed it by down-sampling to 16 kHz (using a Kaiser window filter to avoid aliasing). During the analysis we also investigated how averaging consecutive acoustic features over the following longer time periods affected our results: 1.92, 2.88, 3.84, 4.80, 5.76, 6.72, 7.68, 8.64, 9.60, 29.76, 59.52 and 299.52 s.

### Predictions of species occurrence

For each species we split point counts into two groups; one where the target species was present (*pres*) and the other where it was absent (*abs*). We fit a Dirichlet-process Gaussian mixture model (DP-GMM) to acoustic features from each group to obtain the probability density functions *p*_*pres*_ and *p*_*abs*_ (Blei and Jordan, 2006), using an upper bound of 500 components and diagonal covariance matrices. Other hyperparameters were left as default using the scikit-learn *BayesianGaussianMixture* implementation. For each 20-minute audio recording, we first split the audio into *N* non-overlapping 0.96 s segments. We defined the set *S* of acoustic feature vectors derived from each segment as, *S = {X*_*1*_, *X*_*1*,_ *… X*_*N*_*}*. When using features on longer timescales than 0.96 s, we averaged consecutive members of *S* using non-overlapping windows. For each feature *X*_*i*_ we calculated a likelihood ratio, *L*_*i*_ *= log(p*_*pres*_*(X*_*i*_*)) – log(p*_*abs*_*(X*_*i*_)), allowing us to define a new set, *S*_*L*_ *= {L*_*1*_, *L*_*2*_, *… L*_*N*_*}*. To obtain an overall classification confidence indicating the probability of the species being present in the full 20-minute recording, we looked at four properties of *S*_*L*_; (i) *λ*_*1*_ *= max(S*_*L*_), (ii) *λ*_*2*_ *= min(S*_*L*_), (iii) *λ*_*3*_ *= mean(S*_*L*_), and (iv) *λ*_*4*_ *= P*_*%*_*(S*_*L*_) (for percentiles 10, 20, 30, 40, 50, 60, 70, 80, and 90). We found that the 60^th^ percentile metric, *λ*_*4*_*= P*_*60*_*(S*_*L*_), provided the most accurate predictions, and therefore report results only for this definition of classification confidence (Supp. Fig. S1). Henceforth *λ* will be used to refer to *λ*_*4*_.

To assess the extent to which soundscapes can predict species occurrence we performed an eleven-fold cross-validation classification task for each species. In each fold, data from ten sites were used as a training set (to fit *p*_*pres*_ and *p*_*abs*_), and data from the remaining eleventh site was used as a test set to assess the model’s accuracy. In this way we ensured that we did not report artificially high accuracies by overfitting to site specific soundscapes, but learned generalisable acoustic characteristics that indicated species presence in previously unseen locations. We measured the ability of *λ* to classify a species as present in a point count using the area under the receiver operating characteristic curve (AUC) metric. Mean AUC was calculated for each species across all 11 folds.

For each species we generated null distributions of AUC values to calculate statistical significance of predictions. We used acoustic features at the 2.88s timescale, as these features maximised mean AUC across all species (Supp. Fig. S1). We randomly shuffled classification confidence scores (*λ*) 100 times within each of the 11 folds, and measured AUC using the unshuffled occurrence labels. 100 null mean AUC values were obtained by averaging across the 11 folds, and we used a threshold of *p ≤ 0*.*05* to determine statistical significance.

We performed a similar eleven-fold cross-validation classification task using above-ground biomass data, to compare the predictive power of the two data sources. In each fold, we identified the site in the training set with AGB most similar to the site in the test set. Then, to predict species occurrence in each 20-minute point count, we used the mean species occurrence from point counts at the same time of day from the previously identified similar site.

### Analysis of performance across species

To quantify how temporally structured occurrence patterns were for each species, we formulated a contingency table with species occurrence as one variable and hour of day as the other (using the ground truth point count data). On this contingency table we calculated a χ^2^ statistic. We then calculated Pearson’s correlation coefficient, *ρ*, between the χ^2^ statistic and AUC across all 39 species to test whether accuracy of our predictions was correlated with how temporally structured each species’ occurrence patterns were. We also calculated Pearson’s correlation coefficient between the total number of point counts in which each species was found and AUC to investigate whether rarity of species had an effect on accuracy of predictions. In both cases p-values were obtained analytically.

## Results

### Soundscapes are highly indicative of species occurrence

We were able to predict species occurrence from soundscape recordings for four of the seven non-threatened herpetofaunal species, all 27 non-threatened avifaunal species, and three of the five threatened avifaunal species (*p≤0*.*05*, Fig. 1a). Mean AUC across all species was highest using features on the 2.88 s timescale (Supp. Fig. S1), although the most accurate classifications for a single species was found for the Bold-striped Tit-babbler *Macronus bornensis* when using 0.96 s per feature (0.87 AUC). Variation in AUC between species was larger than the variation for a given species across different timescales of features. Even with features averaged over almost five minutes, we were able to predict species occurrence from soundscapes with AUCs of up to 0.82 (Sooty-capped Babbler *Malacopteron affine*). Spectrograms (Supp. Fig. S2) confirm the intuition that we did not learn to identify species vocalisations, but rather the model learned indicative characteristics of the soundscape that played out over longer timescales than any single species call. From herein we will only consider results using acoustic features at the optimal 2.88 s timescale.

**Figure 1:**
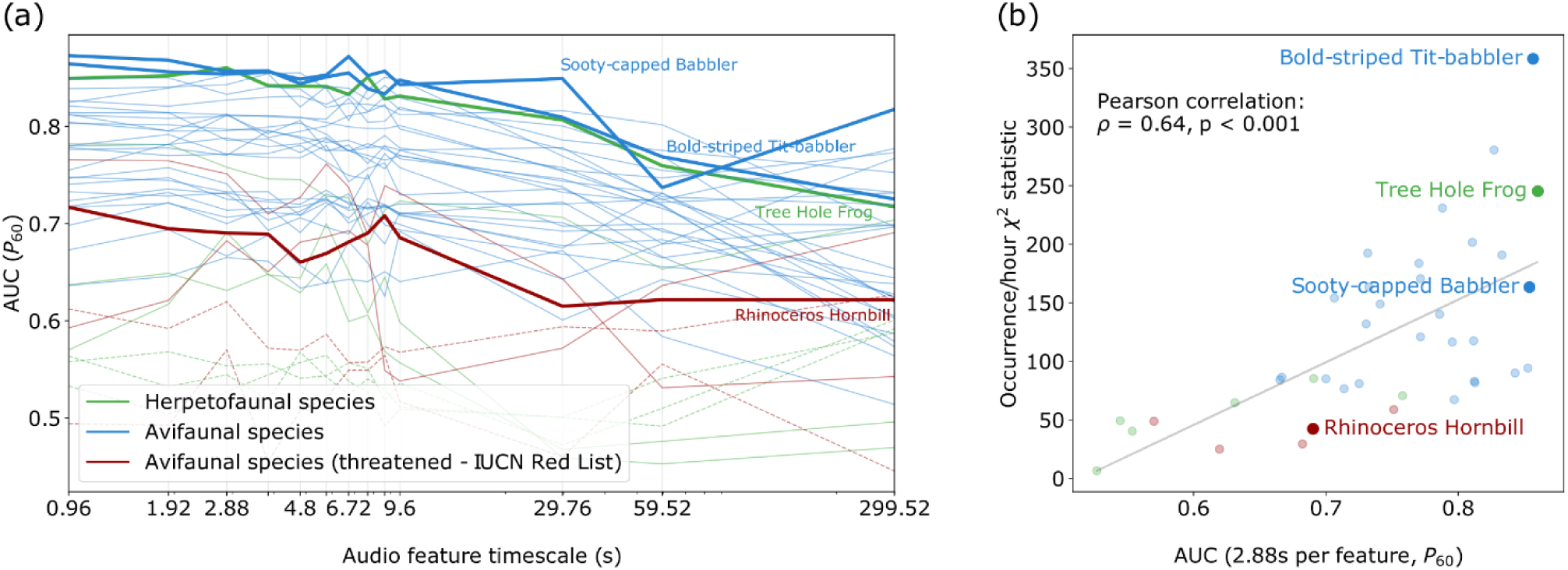
Soundscape features reliably predict species occurrence. We measured how predictive soundscapes were of species presence across 27 non-threatened avifaunal species (blue), five threatened avifaunal species (brown), and seven non-threatened herpetofaunal species (green). **(a)** We found soundscapes features across a wide range of timescales were predictive of species occurrence for 34 species (dotted lines indicate species for which p>0.05). **(b)** The accuracy of occurrence predictions was significantly correlated with a χ^2^ statistic measuring how correlated hour of day was with species occurrence (p<0.001). In both panels we highlight results from four indicative taxa chosen to reflect the variety of species included in this study.

AUC was correlated with total number of encounters of the species across all point counts (Pearson correlation; *ρ=0*.*31, p=0*.*05*). This explains the lower performance for the five rarer Red List threatened avifaunal species compared to the other 27 species (T-test on AUCs; *p<0*.*001*). Nevertheless, occurrence was still predicted with accuracies better than chance (*p≤0*.*05*) for three threatened avifaunal species; the Black Hornbill *Anthracoceros malayanus* (0.69 AUC, n = 15), the Rhinoceros Hornbill *Buceros rhinoceros* (0.69 AUC, n = 34) and the Short-toed Coucal *Centropus rectunguis* (0.75 AUC, n = 23).

We also found that higher AUCs were attained when species were consistently encountered at the same hours of the day (Fig. 1b, Pearson correlation; *ρ=0*.*64, p<0*.*001*). Non-threatened avifaunal species had more temporally structured occurrence patterns than non-threatened herpetofaunal species (T-test on χ^2^ statistics; *p=0*.*04*), explaining the difference in AUCs between the taxonomic groups (T-test on AUCs; *p<0*.*001*). Nevertheless, AUCs for four of the seven herpetofaunal species were still better than would be expected by chance, and reached up to 0.86 for the Tree Hole Frog *Metaphrynella sundana*.

There was a close relationship between predicted occurrence from soundscape data and the pattern of true occurrence across habitat types and time of day (Fig. 2; Supp. Fig. S3 shows similar visualisations for all 39 species). We found that soundscape classification confidence was higher at the true times at which a species would be present, whether the species was diurnal (Fig. 2a, Yellow-vented Bulbul *Pycnonotus goiavier*), nocturnal (Fig. 2c, Tree Hole Frog), or found only during very specific hours (Fig. 2b Sooty-capped Babbler). We also found that soundscape predictions reflected true observations of species habitat niches. For example, the Sooty-capped Babbler (Fig. 2b) and Tree Hole Frog (Fig. 2c) were commonly found in forest habitats – either logged or inside protected areas – whereas the Yellow-vented Bulbul was found more often in heavily disturbed habitats (salvage logged forest and oil palm). In all three cases, classification confidence derived from soundscape data reflected these habitat partitioning patterns.

**Figure 2:**
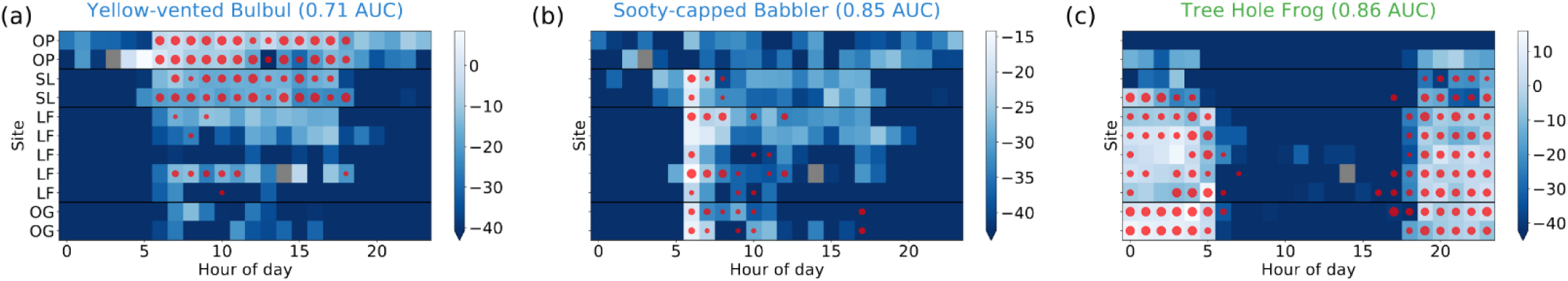
Soundscapes predict occurrence for species with varying habitat and temporal niches. Median classification scores, λ, from dark blue (low) to white (high) are shown for occurrence predictions from soundscapes for three species: **(a)** Yellow-vented Bulbul, **(b)** Sooty-capped Babbler, and **(c)** Tree Hole Frog. Overlaid in red is true occurrence data, where circle sizes indicate how often the species was found at that site and hour during the manual point counts. Sites are ordered by above-ground biomass with low-quality habitats at the top and high-quality habitats at the bottom. (Supp. Fig. S3 provides the same visualisation for all 39 species).

### Soundscapes predict occurrence more accurately than above-ground biomass

We found that soundscape features predicted species occurrence more accurately than a comparison model based on above-ground biomass (AGB) data, a metric often used as a proxy for tropical forest habitat quality (Fig. 3, paired T-test on AUCs; *p<0*.*001*). The soundscape-based model produced increased AUCs for 31 of the 39 species surveyed, including for all five threatened avifaunal species. Mean accuracy of occurrence predictions for the non-threatened avifaunal group was increased by 0.08 AUC, for the threatened avifaunal group by 0.06 AUC, and for the non-threatened herpetofaunal group by 0.03 AUC. This followed the trends noted earlier, as groups of species which were common, or exhibited strong temporal occurrence patterns benefited the most from the soundscape based approach. Per species there was a mean percentage increase in AUC of 10 % across all 39 species surveyed.

**Figure 3:**
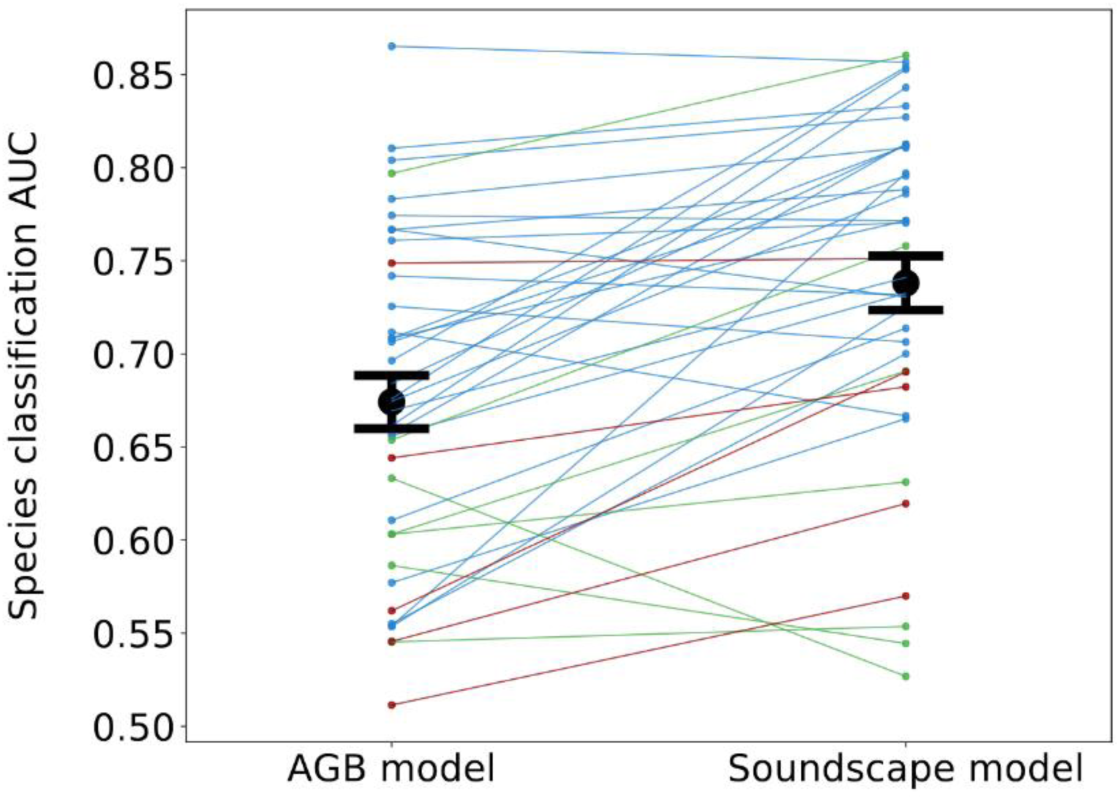
Soundscape features are a better indicator of species occurrence than above-ground biomass (AGB). We compared occurrence predictions using soundscapes to a comparison model using AGB data. Lines connect AUC metrics for the same species, with threatened avifaunal species in brown, non-threatened avifaunal species in blue, and non-threatened herpetofaunal species in green. In black is the mean and standard error for AUC across all 39 species for each model.

## Discussion

We investigated whether soundscapes could indicate the occurrence patterns of 39 species across two taxonomic groups. Our results demonstrate this is indeed feasible, and that the most accurate indications can be obtained for species for which we have the most data (the most common species) and those with strong temporal occurrence patterns. This meant that our approach is less suited to the monitoring of endangered species, although we were able to successfully model occurrence for three species based on very sparce occurrence data of just 15, 23, and 34 observations across 790 point counts. Nevertheless, species listed as vulnerable or endangered by the IUCN Red List are not the only ones of conservation interest. Species which are particularly good indicators of habitat quality, those that have a disproportionate ecological impact on their environment, or those that fulfil important economic functions are often referred to as “keystone species” (Mills et al., 1993). Whilst these species are sometimes also endangered, this is not always the case. For example, within the “non-threatened” species, we had the Rough Guardian Frog *Limnonectes finchi*, a species only ever found close to suitable water sources (Inger and Voris, 1988). We also had the White Crowned Shama *Copsychus stricklandii* which due to their unique singing ability is threatened by a high rate of unsustainable trade in South East Asia (Leupen et al., 2018). We were able to predict occurrence for both of these species accurately. Using soundscapes to assess habitat suitability for species like these would therefore be a promising route to take, with valuable conservation data to be gained.

We found that our model was most accurate when using short timescale acoustic features. This may simply be a matter of resolution – with longer timescale features the details of how soundscapes move between different modes are lost. The average of shorter features over these long time periods will therefore only provide a crude overview of the overall soundscape, leading to less accurate predictions of occurrence. Nonetheless, there was still significant predictive information contained within long timescale features, indicating that a coarse acoustic overview is often all that is required.

Our model learned to identify soundscape features that were uniquely found when the species of interest was present. In this study, we found that this did not correspond to species vocalisations, but sounds typical to the habitat type, or time of day that the species was likely to be found. This was probably due to the low number of positive samples we had for each species, together with the high overall temporal and spatial variability of soundscapes across all of our audio recordings. Whilst all species surveyed in this study were vocal, foregoing a reliance on vocalisations means that our approach can be used to explicitly predict the occurrence of completely silent species. Equally tantalisingly, there is a possibility that with a less heterogenous, larger dataset a similar approach to ours may enable identification of species vocalisations in an unsupervised manner. This would occur if the predominant distinguishing acoustic features between present and absent samples was the sound of the species vocalising. Automatically extracting vocalisations from passive recordings *in situ* may even allow us to discover calls and behaviours that cannot be reproduced with the same species in a more controlled environment.

Other types of data, beyond audio, can be used to predict species occurrence at a given place and time. AGB is a habitat quality indicator used for many species across tropical forest degradation gradients, and it has been used extensively at the field site we surveyed (e.g., Brant et al., 2016; Luke et al., 2017; Riutta et al., 2018). In this study, however, we showed that soundscapes were in fact better predictors of species occurrence for 31 of the 39 species surveyed. Furthermore, manual field surveys to collect AGB data are cumbersome (Pfeifer et al., 2015), and when data from planes or satellites are used the costs can be prohibitively high (Lefsky et al., 2002; Popescu et al., 2011). By contrast, our audio recording protocol only involved using an inexpensive handheld recorder deployed to gather a 24 hour acoustic record per site. Recordings of this type could be made rapidly from a large number of sites, providing wide coverage with minimal capital outlay.

The link between habitat suitability and species occurrence data is clear – species are more likely to be found in habitats that are able to sustainably support their needs (Hirzel et al., 2006). Thus, by showing that occurrence for a wide range of species can be accurately predicted by soundscapes, this opens up a new avenue for assessing habitat suitability from audio data. One use-case may be in assisting the identification of areas of high conservation value within agricultural landscapes, as required by certification agencies such as the Roundtable for Sustainable Palm Oil (Brown et al., 2013). Additionally, as collaborative eco-acoustic datasets continue grow (Baker et al., 2015), we may be able to harness soundscape data to produce large-scale habitat suitability maps, and identify those species that are most at risk from global pressures such as climate change (Walther et al., 2002).

## Conclusion

In this study we have demonstrated that soundscapes can be used to predict species occurrence across a wide range of species in tropical forests. We found that the most accurate predictions could be made for common species with strong temporal occurrence patterns, including for species of specific conservation concern. Using a comparison model, we found that soundscapes were able to predict occurrence more accurately than above-ground biomass, a widely used indicator of habitat quality across forest degradation gradients. Our findings indicate a promising new route for audio data to be used as an impactful conservation tool whilst side-stepping many of the scalability issues of existing approaches.

## Author’s contributions

S.S.S., R.M.E., N.S.J., and L.P. contributed to the conceptualisation, development of analysis methods and final implementation of this study. S.S.S., J.S., A.S., and N.Z collected the field data. S.S.S., R.M.E., N.S.J., and L.P. led the manuscript writing process, with input provided from all authors.

### Acknowledgements

Thanks to Sahil Loomba and Henry Bernard for their contributions to the study, and to the staff at the SAFE Project and SEARRP for logistical support in data collection. This project was supported by funding from WWF (Biome Health Project), Sime Darby Foundation (SAFE Project), NERC (NE/K007270/1, N.S.J.), and EPSRC (EP/R511547/1, S.S.S., EP/N014529/1, N.S.J.). Data was collected from Malaysia under an SaBC permit granted to S.S.S. (JKM/MBS.1000-2/2 JLD.8 (63)).

## Data availability

Raw data from the point counts can be found at https://doi.org/10.5281/zenodo.3265711 (Sethi et al., 2019). Processed audio and field data are stored at https://zenodo.org/record/4048019#.X2X2Gu17kuU, and code to reproduce results and figures is at https://github.com/sarabsethi/sscape_spec_occ_preds_sethi2020.

## Supplementary Information

**Table S1:** We surveyed 39 species across two taxonomic groups. Here we provide common names, Latin binomials, IUCN Red List status, and total number of occurrences across all point counts for each species we consider in this study. Red List acronyms are as follows: LC = least concern, NT = near threatened, VU = vulnerable, CR = critically endangered. We defined threatened species to be in either VU or CR categories. Non-threatened herpetofaunal species are in green, non-threatened avifaunal species in blue, and threatened avifaunal species are in brown.

| Species common name | Latin binomial | Red List status | N occurrences |
| --- | --- | --- | --- |
| Cricket Frog | *Hylarana nicobariensis* | LC | 121 |
| Four-lined Tree Frog | *Polypedates leucomystax* | LC | 75 |
| Grass Frog | *Fejervarya limnocharis* | LC | 63 |
| House Gecko | *Hemidactylus frenatus* | LC | 117 |
| Rough Guardian Frog | *Limnonectes finchi* | LC | 78 |
| Smith's Giant Gecko | *Gekko smithii* | LC | 71 |
| Tree Hole Frog | *Metaphrynella sundana* | LC | 211 |
| Ashy Tailorbird | *Orthotomus ruficeps* | LC | 143 |
| Asian Red-eyed Bulbul | *Pycnonotus brunneus* | LC | 62 |
| Black-and-yellow Broadbill | *Eurylaimus ochromalus* | NT | 63 |
| Black-headed Bulbul | *Pycnonotus atriceps* | LC | 126 |
| Black-naped Monarch | *Hypothymis azurea* | LC | 53 |
| Blue-eared Barbet | *Psilopogon duvaucelii* | LC | 82 |
| Bold-striped Tit-babbler | *Macronus bornensis* | LC | 269 |
| Chestnut-backed Scimitar Babbler | *Pomatorhinus montanus* | LC | 66 |
| Chestnut-winged Babbler | *Cyanoderma erythropterum* | LC | 182 |
| Common Emerald Dove | *Chalcophaps indica* | LC | 55 |
| Dark-necked Tailorbird | *Orthotomus atrogularis* | LC | 68 |
| Fluffy-backed Tit-babbler | *Macronus ptilosus* | NT | 134 |
| Great Argus | *Argusianus argus* | NT | 90 |
| Greater Coucal | *Centropus sinensis* | LC | 139 |
| Lesser Coucal | *Centropus bengalensis* | LC | 74 |
| Little Spiderhunter | *Arachnothera longirostra* | LC | 170 |
| Malaysian Pied Fantail | *Rhipidura javanica* | LC | 119 |
| Plaintive Cuckoo | *Cacomantis merulinus* | LC | 94 |
| Rufous-tailed Tailorbird | *Orthotomus sericeus* | LC | 164 |
| Short-tailed Babbler | *Pellorneum malaccense* | NT | 69 |
| Slender-billed Crow | *Corvus enca* | LC | 113 |
| Sooty-capped Babbler | *Malacopteron affine* | NT | 55 |
| Spectacled Bulbul | *Pycnonotus erythrophthalmos* | LC | 158 |
| White-crowned Shama | *Copsychus stricklandii* | LC | 106 |
| Yellow-bellied Prinia | *Prinia flaviventris* | LC | 179 |
| Yellow-rumped Flowerpecker | *Prionochilus xanthopygius* | LC | 56 |
| Yellow-vented Bulbul | *Pycnonotus goiavier* | LC | 143 |
| Black Hornbill | *Anthraceroceros malayanus* | VU | 15 |
| Chestnut-necklaced Partridge | *Arborophila charltonii* | VU | 21 |
| Helmeted Hornbill | *Rhinoplax vigil* | CR | 16 |
| Rhinoceros Hornbill | *Buceros rhinoceros* | VU | 34 |
| Short-toed Coucal | *Centropus rectunguis* | VU | 23 |

**Figure S2:**
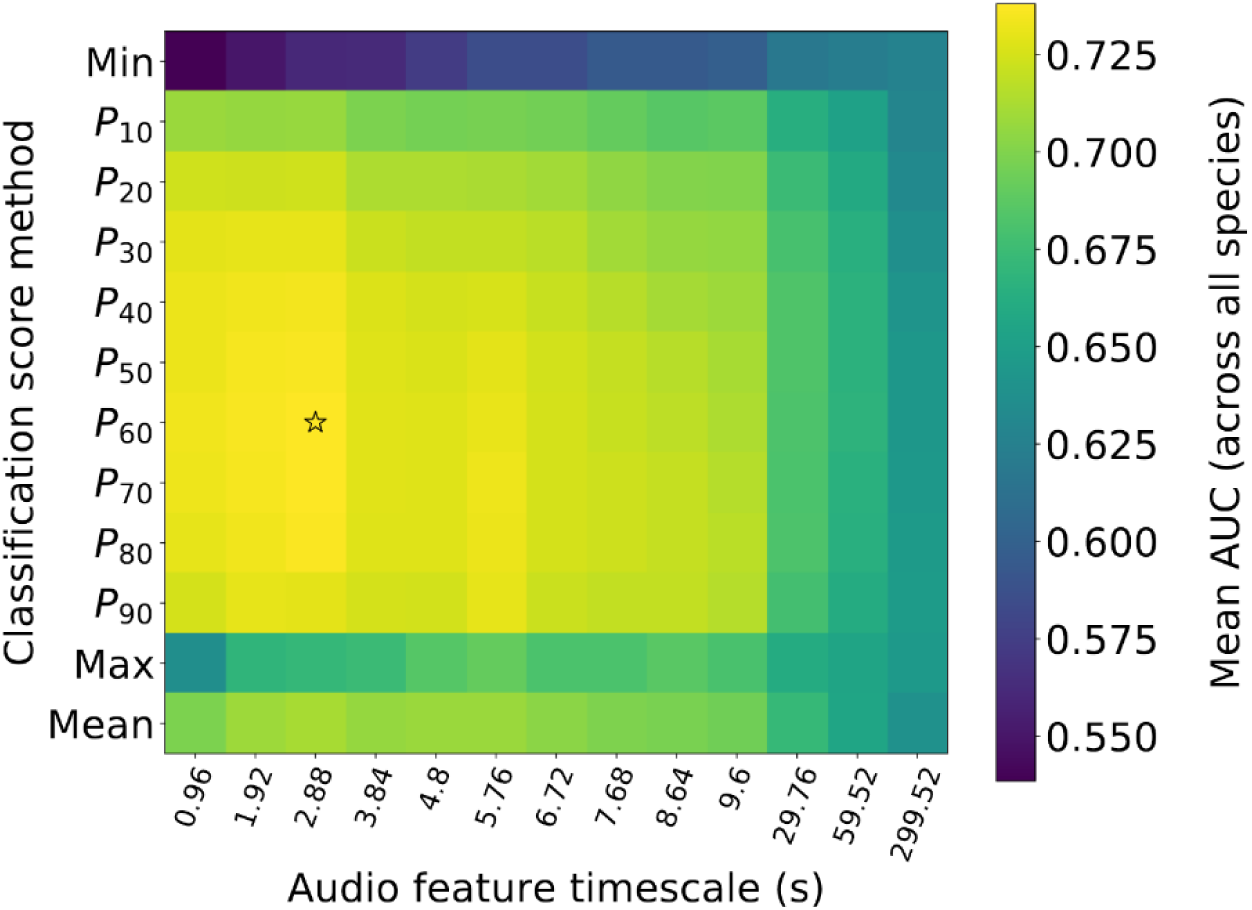
A grid search reveals the optimal parameters for classification using soundscape features. We performed a grid search across acoustic feature timescales and method of deriving classification confidence, λ. For each combination, we calculated mean AUC of classifications across all 39 species. We found acoustic features on the 2.88s timescale with the P_60_ (60^th^ percentile) metric provided the most accurate classifications.

**Figure S3:**
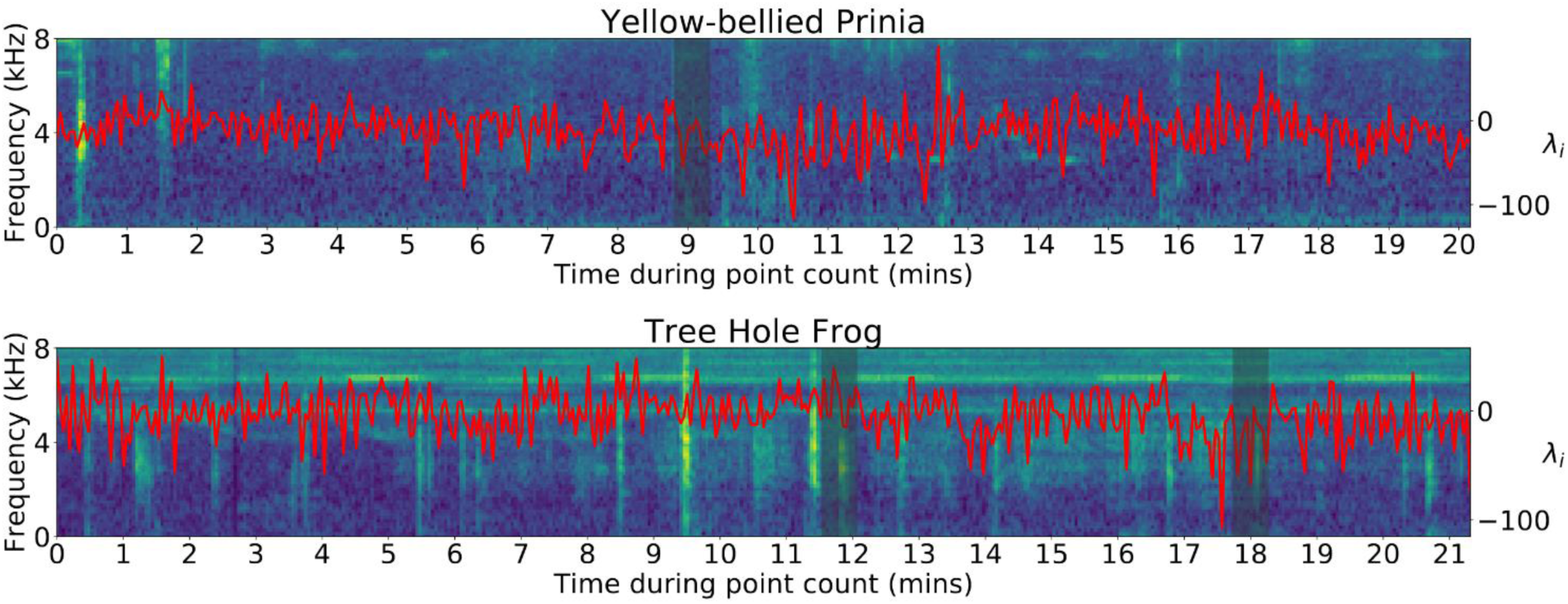
Classifications are not based on species vocalisations. We visualise per feature classification confidence (λ_i_, red) together with a spectrogram of the audio data from one point count each for the Yellow-bellied Prinia and Tree Hole Frog. Shaded are regions during which each of the species is vocalising. Classification confidence does not increase significantly during these periods, indicating that predictions of occurrence are not based on exact species vocalisations, but other components of the overall soundscape that indicate species presence.

**Figure S4:**
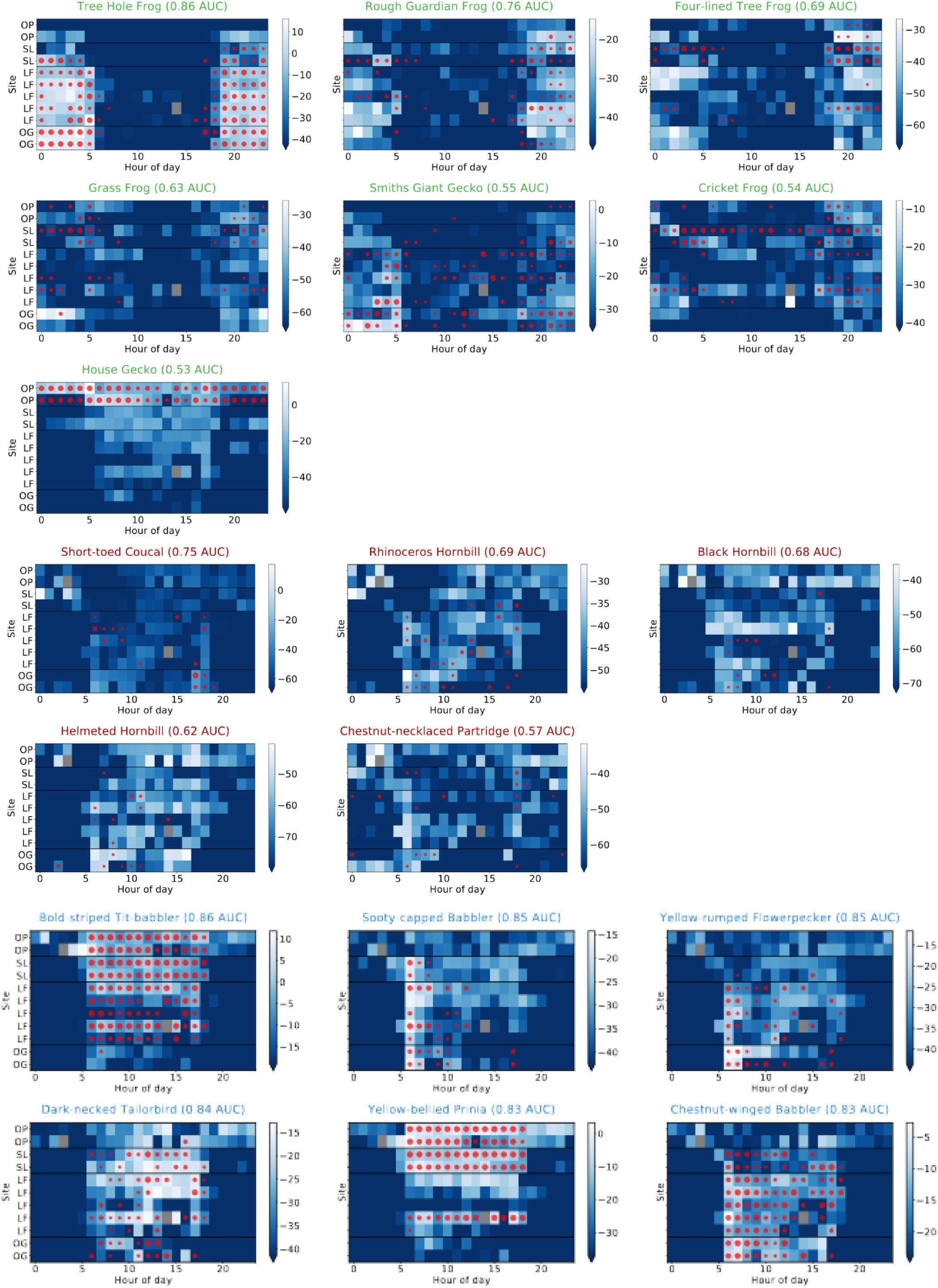
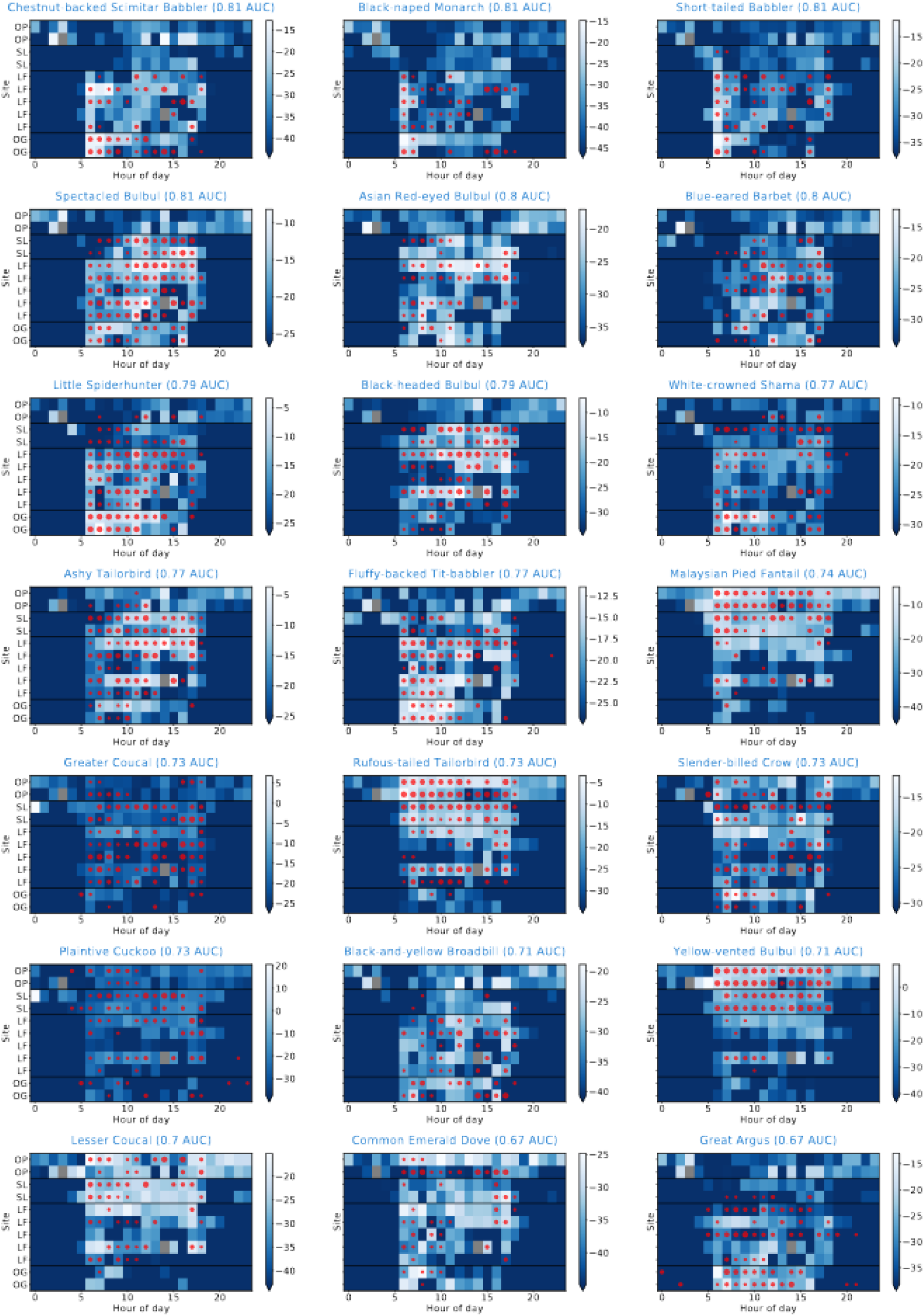
Classification confidence from soundscape data reflects true occurrence data for many species. Median classification scores, λ, from blue (low) to white (high) are shown for occurrence predictions from soundscapes for all 39 species tested. Overlaid in red is true occurrence data, where circle sizes indicate regularity with which the species was found at that site and hour.

